# Triatomine host preference in an experimental cafeteria: the impact of *Trypanosoma cruzi* infections

**DOI:** 10.64898/2026.01.12.698978

**Authors:** Jordan Salomon, Jeffery K. Tomberlin, Keswick C. Killets, Dorcas Abiara, Hojun Song, Gabriel L. Hamer, Sarah A. Hamer

## Abstract

**Background:** Hematophagous triatomine vectors of the Chagas disease parasite, *Trypanosoma cruzi*, have a wide bloodmeal host breadth yet mammals are the primary competent reservoirs for *T. cruzi*. Accordingly, the transmission of *T. cruzi* is dependent on triatomines encountering mammalian hosts. We hypothesized *T. cruzi*-infected triatomines exhibit a preference for mammalian versus non-mammalian hosts, as compared to uninfected triatomines.

**Methodology/Principal Findings:** We used a dual-choice olfactometer to present individual *T. cruzi-*infected and uninfected *Triatoma gerstaeckeri*- an important North American triatomine species- to pairwise combinations of no host, dog, human, or chicken. We found that uninfected *T. gerstaeckeri* preferred dogs over chickens and humans, but *T. cruzi-*infected *T. gerstaeckeri* had no host preference. Additionally, *T. cruzi-*infected *T. gerstaeckeri* took less time to begin advancing towards hosts (average of 522 s ± 577 s vs 801 s ± 704 s) and spent more time on both sides of the olfactometer compared to uninfected insects As a comparative group,we tested *Rhodnius prolixus*- a potent South American vector- and found no statistical preference among hosts, yet they advanced towards hosts more often (63.2% vs 47.1%) and faster compared to *T. gerstaeckeri*.

**Conclusions/Significance:** The increased locomotion of *R. prolixus* and *T. cruzi*-infected *T. gerstaeckeri* toward hosts may promote increased transmission of *T. cruzi*. Dogs are a preferred host for *T. gerstaeckeri* and may be considered in future host-targeted interventions for triatomine control.

**Author Summary:** Vectors are often generalists or opportunists with a wide range of available hosts, yet the parasites they transmit may have a narrower host range. It is in the parasite’s best interest for the vectors they infect to interact with competent hosts, thereby introducing the setting for parasite manipulation of vectors. We aimed to understand the foraging behaviors of vectors and how those behaviors may be different when the vector is infected with a parasite. We studied the triatomine (‘kissing bug’) insect vectors of the Chagas disease parasite, *Trypanosoma cruzi*, using one of North America’s most important vector species in comparison to a potent South American vector species. We tested if parasite-infected triatomines exhibited a preference toward vertebrate hosts that are also competent reservoirs for the *T. cruzi* (i.e., mammals). We used an experimental cafeteria, in which individual vectors were introduced to a dual-choice olfactometer and allowed to walk toward hosts including restrained humans, dogs, or chickens. We found differences in apparent host preference between species of vectors and between infected vs. uninfected vectors. Uninfected vectors displayed a host preference toward dogs, yet this preference was not significant among infected vectors. Uninfected vectors had higher frequency of locomotion in the arena, suggesting the opportunity for more vector-host contact.

## Introduction

Most vector-borne zoonotic diseases are caused by parasites transmitted by vector species that feed on multiple host species. Host availability and host preference therefore sculpt feeding patterns in nature. Parasite manipulation of vector foraging behaviors to increase parasite fitness (i.e. transmission) is a widespread phenomenon in vector-borne zoonotic disease systems [1]. For instance, parasites can manipulate the chemical composition of a vector’s saliva to increase blood-feeding, or influence vector movement in the presence of hosts to increase predation of the vector [2–4]. In some systems, hosts infected with parasites are more attractive to vectors [5,6]. Interactions among vectors, parasites, and hosts- and in particular vector foraging behaviors – are a key driving factor of human and animal risk of infection.

Observed vector-feeding patterns may or may not reflect vector-host preference [7]. Vector-feeding patterns on hosts are driven by both intrinsic (i.e. genetic) and extrinsic (i.e. host availability) factors, which can be inferred from data which includes direct removal of vector from the host or analysis of vector bloodmeals to identify the species upon which the vector previously fed. In contrast, host preference is driven by only intrinsic factors and experimental tests for preference require a decision-based (i.e. a choice) assay by providing the vector multiple host options. With the desire to learn which host species are most critical in supporting vector-borne parasite transmission that leads to human disease risk, researchers may first identify vector-host feeding patterns through surveillance and next experimentally test for preference among key host taxa, yet such investigations should reflect that parasite manipulation may drive vector foraging behaviors to increase opportunities for transmission.

*Trypanosoma cruzi*, the agent of Chagas disease, is vectored by triatomine insects (Hemiptera: Reduviidae: Triatominae). Approximately 8 million humans are diagnosed with Chagas disease, a neglected tropical disease that is endemic to South America, Central America, Mexico and the southern United States—regions where the triatomine vector and the parasite occur [8,9]. There are multiple routes of transmission, but the most common are via contact with triatomine feces contaminated with *T. cruzi* or the ingestion of an infected triatomine. Wildlife, domesticated animals, and humans are all important hosts for the parasite and vectors. As a neglected tropical disease [10], there is no vaccine, diagnostic tests are imperfect, and treatments are limited. There is a spectrum of medical outcomes of infection in humans and dogs [8,11], which can range from heart disease, gastrointestinal diseases, and even death [12].

Despite the ability of triatomines to survive on low-risk nutritional sources, such as plants [13,14] and other insects [15], they predominantly feed on blood of vertebrates. Vector-feeding pattern studies show triatomines are generalists utilizing many different vertebrate bloodmeals; but within domestic habitats, across triatomine species a high proportion of bloodmeals are consistently recorded from three key species: chickens, dogs, and humans [8,16–18]. Chickens are refractory for *T. cruzi* as their blood lyses the parasite preventing parasitemia [19]. In contrast, dogs and humans are competent for *T. cruzi* and develop parasitemia capable of infecting feeding triatomines [8,11]. Although both dogs and humans maintain *T. cruzi* in their bloodstream for only about a month after exposure [11], high triatomine densities can result in repeated *T. cruzi* exposure, prolonging the duration of parasitemia [12]. Consequently, triatomine feeding and host contact dictates the reproductive success of *T. cruzi* and ultimately the risk of infection to susceptible hosts.

Like other arthropod vectors, triatomines are sensitive to the physical stimuli of light and temperature, in addition to the chemical stimuli of humidity, CO_2_, and some odorants [20]. These stimuli collectively attract triatomines toward a vertebrate bloodmeal [20], but they vary between hosts and could potentially elicit different attractiveness. Furthermore, the triatomines infected with *Trypanosoma* spp. have been shown to have an increase in locomotion during daytime [21], expression of a gene responsible for locomotion [22], sensilla abundance on antennae [23], and response time to host olfactory cues [24], suggesting the parasite’s capacity to manipulate foraging behavior. However, whether *T. cruzi* can manipulate triatomine preference for hosts with variable reservoir competency for the parasite is currently unknown.

In this study, we tested if parasite infection of vectors is associated with altered foraging behaviors including vector-host preferences. We used a dual-choice olfactometer to compare between *T. cruzi-*infected and uninfected triatomines when presented with frequently utilized hosts that vary in competence for *T. cruzi* (i.e., chickens, dogs, or humans). We compared the vector-host preferences of two different species of triatomines, a species endemic to Texas (*Triatoma gerstaeckeri*) [9,25–29] and a South American species commonly found in domestic settings (*Rhodnius prolixus*). An understanding of triatomine foraging behaviors regarding host feeding preferences and how it differs depending on the parasite infection would provide critical information for future investigations of parasite-host coevolutionary mechanisms; in addition to the application of specific targets for more efficient deployment of management strategies [30–35].

## Methods

### Source of dog and human volunteers

Dog and human volunteers were recruited for the study through emails among Texas A&M University and dog training programs in College Station, Texas, USA (IRB2021-0945F, IACUC 2022-0001 CA). Inclusion criteria for all hosts to be enrolled in the study necessitated that they were comfortable with sitting in a dark box enclosed for up to an hour. Exclusion criteria for dog volunteers included use of a topical insecticide or testing positive for *T. cruzi* antibodies (see Supplemental Materials for details). Using fear free, positive reinforcement and shaping learning techniques [36–38], volunteer dogs were acclimated to the boxes one day prior to the trial. This acclimation was done in a series of successive approximation where each motion of going into the box and closing it was followed by a treat reward if the dog was showing no signs of distress (i.e., barking, crying, avoiding, scratching). Dogs moved forward only if they showed no distress with this acclimatization process. On the day of the study, humans were asked to not use any fragrances (i.e., perfume or cologne) or topical insect repellants. Height and weight were recorded for both humans and dogs, and approximate age was also recorded for dogs.

### Source of chickens

Female Hy-Line W36 chickens were owned and housed by the Texas A&M University Poultry Science Center where they were provided a standard layer diet and water *ad libitum* (IACUC 2021-0109). Chickens used in the study were not exposed to insecticides or repellents. Chickens were transported in a cat carrier (44.45 cm long, 30.48 cm wide, and 19.05 cm tall) from the chicken farm to the isolation building containing the olfactometer for trials and remained inside the carrier for the duration of the trials.

### Source of triatomines

Triatomines were sourced from a colony maintained at Texas A&M University in a USDA-APHIS PPQ-approved arthropod containment level 2 (ACL2) quarantine facility. Offspring (F1 or F2) from wild caught *Triatoma gerstaeckeri* from Texas were used in the trials. The *Rhodnius prolixus* colony was originally sourced from Centers of Disease Control and Prevention from a colony developed from insects collected in Colombia (NR-44077, BEI Resources, Manassas, Virginia, US) and was maintained at the Texas A&M University insectary for 5 years prior to use in these trials. In the quarantine facility, triatomines were maintained at 27–33 °C, 30–60% relative humidity, and a 12-hr photoperiod with a weekly offering of defibrinated rabbit (*Oryctolagus cuniculus domestics*) blood (HemoStat Laboratories, Dixon, California, US) warmed to 37°C and secured with a parafilm membrane using Hemotek membrane feeders (Hemotek Ltd, Lancashire, UK) [39]. The 4^th^ and 5^th^ instar nymphs were removed from their original colony and placed in an isolated container for starvation prior to trials. Triatomines were starved for a minimum of 13 days prior to their use in a trial, and duration of starvation was included as a variable in statistical models.

### Experiemtal infection of T. gerstaeckeri with T. cruzi

*Triatoma gerstaeckeri* were infected with *T. cruzi* by inoculating defibrinated rabbit blood with live *T. cruzi* epimastigote cultures [40]. Cultures of *T. cruzi* discrete typing unit (DTU) TcI were initially established from the feces of live-wild-caught triatomines that were collected for a community science program [25,29]. The culture media consisted of liver infusion broth (DIFCO Laboratories Inc., San Diego, CA, US), sodium chloride, potassium chloride, monohydrated glucose, Bacto^TM^ tryptose (DIFCO Laboratories Inc.), sodium phosphate dibasic, hemin (Sigma-Aldrich, St. Louis, MO, US), sterile inactivated fetal bovine serum, antibiotics (Penicillin-Streptomycin), and antifungals (Nystatin) [41]. Once inoculated, cultures were maintained in an incubator set to 27° C and were checked regularly for growth of *T. cruzi*. Once *T. cruzi* was in the exponential growth phase, a target concentration of 3 × 10^6^ parasites per mL of culture media was separated and washed prior to adding it to the defibrinated rabbit blood following protocols previously described in [40]. The infected blood was fed to triatomines through the Hemotek apparatus inside a biosafety cabinet. Once triatomines were engorged with the *T. cruzi*-inoculated blood, they were isolated individually into 50-mL conical tubes lined with filter paper.

To verify *T. cruzi* infection of *T. gerstaeckeri* prior to a trial, we tested feces that were collected on filter paper via a *T. cruzi* qPCR protocol two weeks post-infected bloodmeal [42] following DNA extraction (Omega Bio-tek, Norcross, GA, US; Qiagen, Germantown, MD, US). *Triatoma gerstaeckeri* individuals with a qPCR negative fecal spot were fed again blood with *T. cruzi* and were not used in trials until they produced a qPCR-positive fecal spot.

### Olfactometer design

To quantify triatomine host preference, we built a horizontal “T” dual-choice olfactometer, in which a single triatomine had the opportunity to travel from a central location towards a host (Figure 1). Two equally sized host chambers (127 cm by 73.9 cm by 104.14 cm, KINYING Outdoor Storage Shed, Zhejiang, China) were connected by clear PVC pipes (6.35 cm diameter, POWERTEC, Chicago, IL, US) to allow the triatomine to move between host chambers. At the center of these two clear PVC pipes was a 1.5 cm hole for which we were able to measure abiotic outputs (temperature, humidity, and windspeed) from both host chambers periodically throughout the trial. When measurements were not being taken, a rubber stopper was placed in the holes. Air was pulled through the PVC pipes to a clear PVC flexible hose (15.24 cm radius) leading to an external exhaust fan pulling air out of the room to create negative air pressure in the animal isolation room. Before the T-split, the single PVC tube contained a port which allowed the triatomines to be placed into the olfactometer. To create the “T” formation, the fan duct was connected to a clear “Y” PCV connection (15.24 cm length and 6.35 cm diameter, POWERTEC), which was then connected to the “T” coupling (10.16 cm length and 6.35 cm diameter, POWERTEC) to connect to the clear PVC tube which connects both host chambers. The total distance of the piping connecting the two host chambers was 91.44 cm before the pitfall traps.

**Figure 1:**
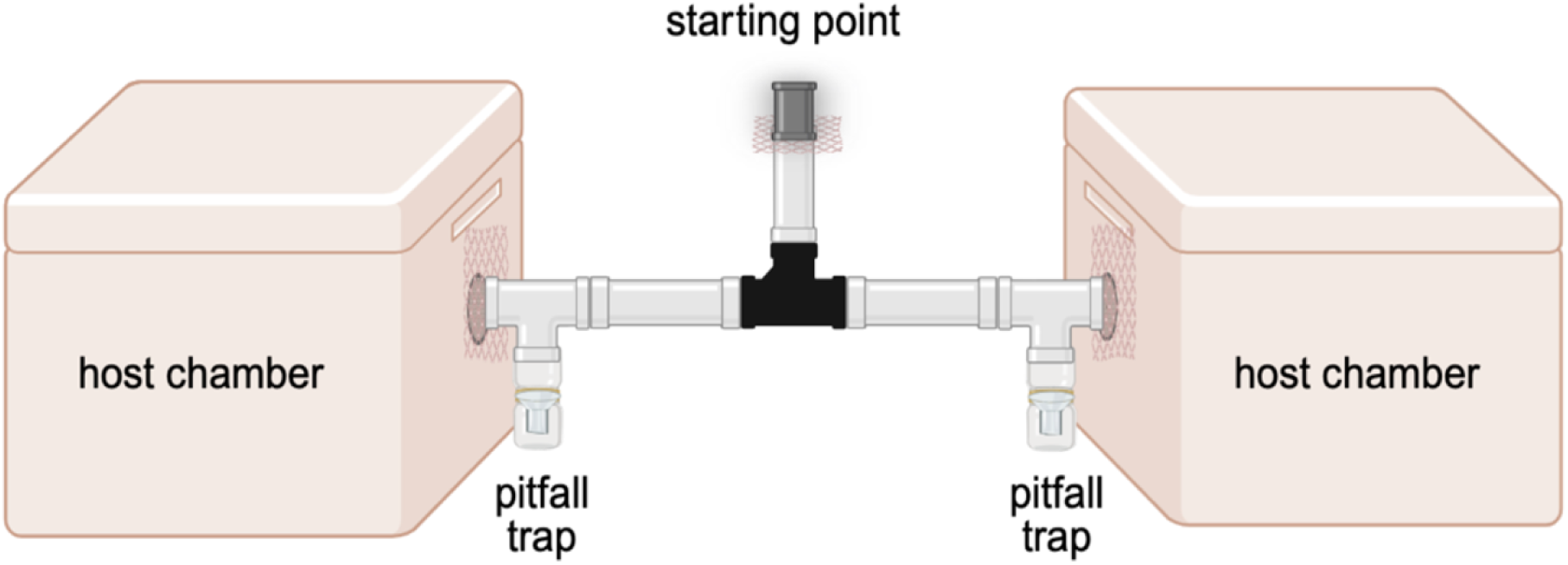
Diagram of the olfactometer with dual-choice “T” design used for the study of triatomine foraging behavior and host choice.

Pitfall traps were made with clear “Y” PCV connections (15.24 cm length and 6.35 cm diameter, POWERTEC) glued to a clear mosquito rearing container. Pitfall traps were connected to the host chambers via rubber couplings with stainless steel clamps (6.35 cm diameter) to a black cone reducer (10.16 cm to 6.35 cm diameter, POWERTEC) and a black rectangular dust hood (10.16 cm diameter, POWERTEC).

To prevent triatomine escape, five cloth mesh squares were secured in between the rubber couplings and between the inline fan duct and where the triatomine is inserted into the olfactometer. Additionally, to prevent triatomine access to hosts, a barrier was created with steel woven mesh (1mm mesh size) to cover the air flow outlet (Figure 1). Within the room of the olfactometer, all air ducts were covered with activated charcoal air filters (Breathe Naturally, Rochester, MN, USA) to filter all air that enters the room.

### Experimental trial designs

Each trial consisted of various pairwise combinations of host or empty chamber (Figure 2-4). We randomized the pairwise combinations and order of trials by generating a list using *“lapply”* in base R package with four options being dog, chicken, human, or empty. Control trials (empty versus empty) were added to the randomized trial list using a random number generator. All trials were conducted in the order of the list with minor variation based on the availability of live subjects.

**Figure 2:**
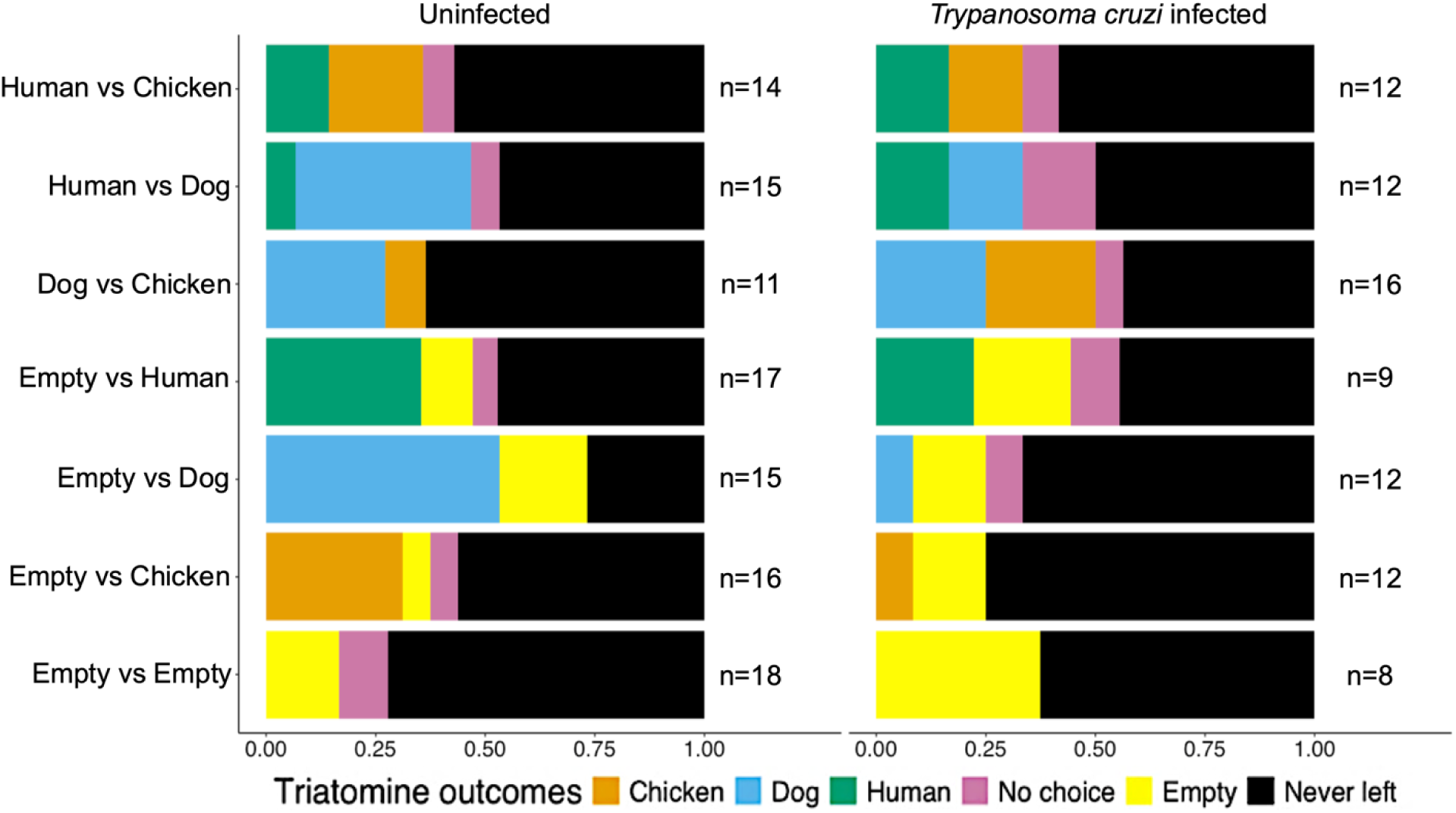
The proportion of trial outcomes among individual *T. gerstaeckeri* uninfected or infected with *T. cruzi*. Each bar represents the trial combinations, and the colors represent a different triatomine outcome for that trial combination. The black numbers to the right of each bar represent the total number of triatomines used for that trial combination.

**Figure 3:**
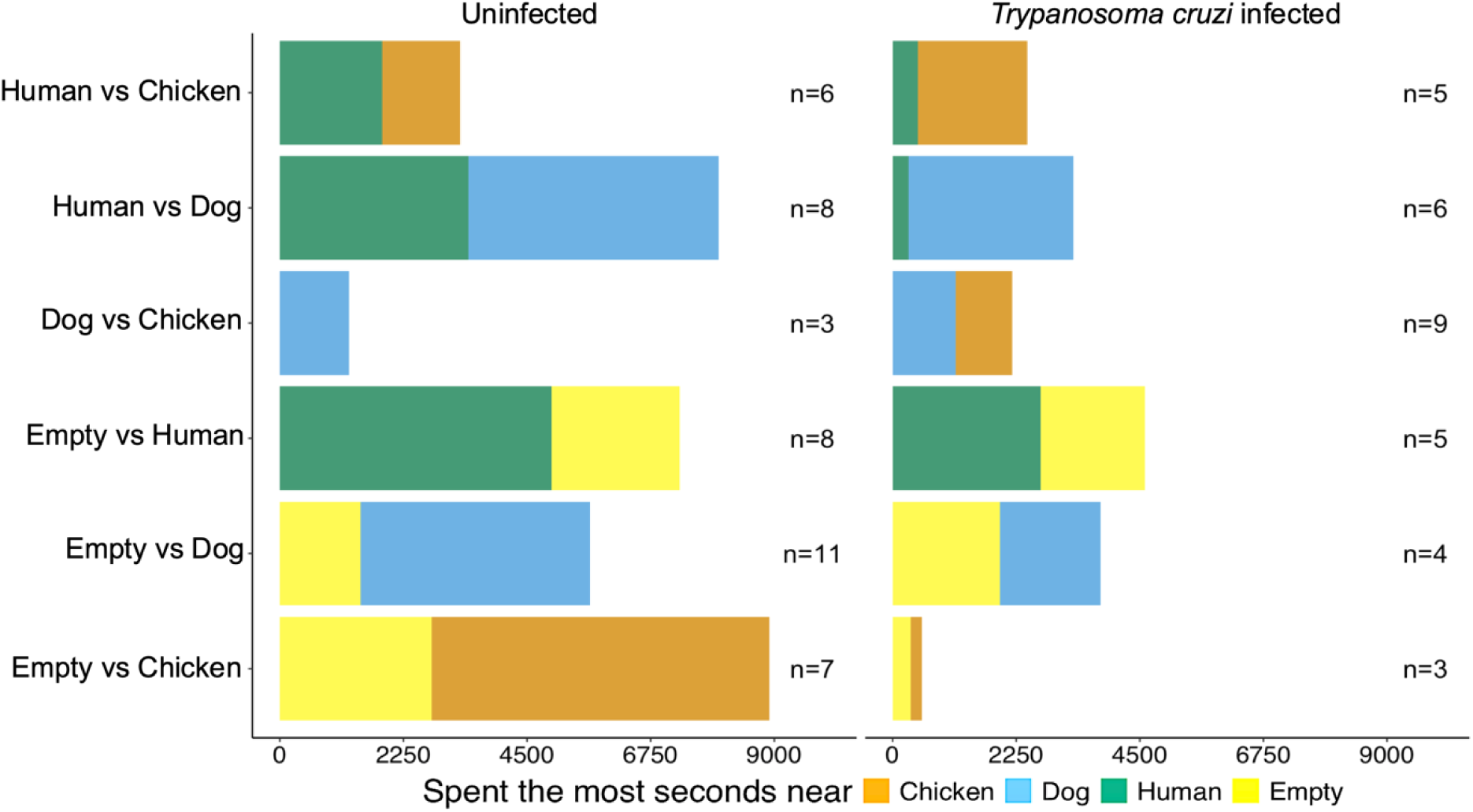
The duration of time in seconds spent by *T. gerstaeckeri* uninfected or infected with *T. cruzi* in the olfactometer advancing towards host options. Each bar represents the trial combinations, and the colors represent a different outcome in which the triatomine spent the most time near a host. The black numbers to the right of each bar represent the total number of triatomines used for that trial combination.

**Figure 4:**
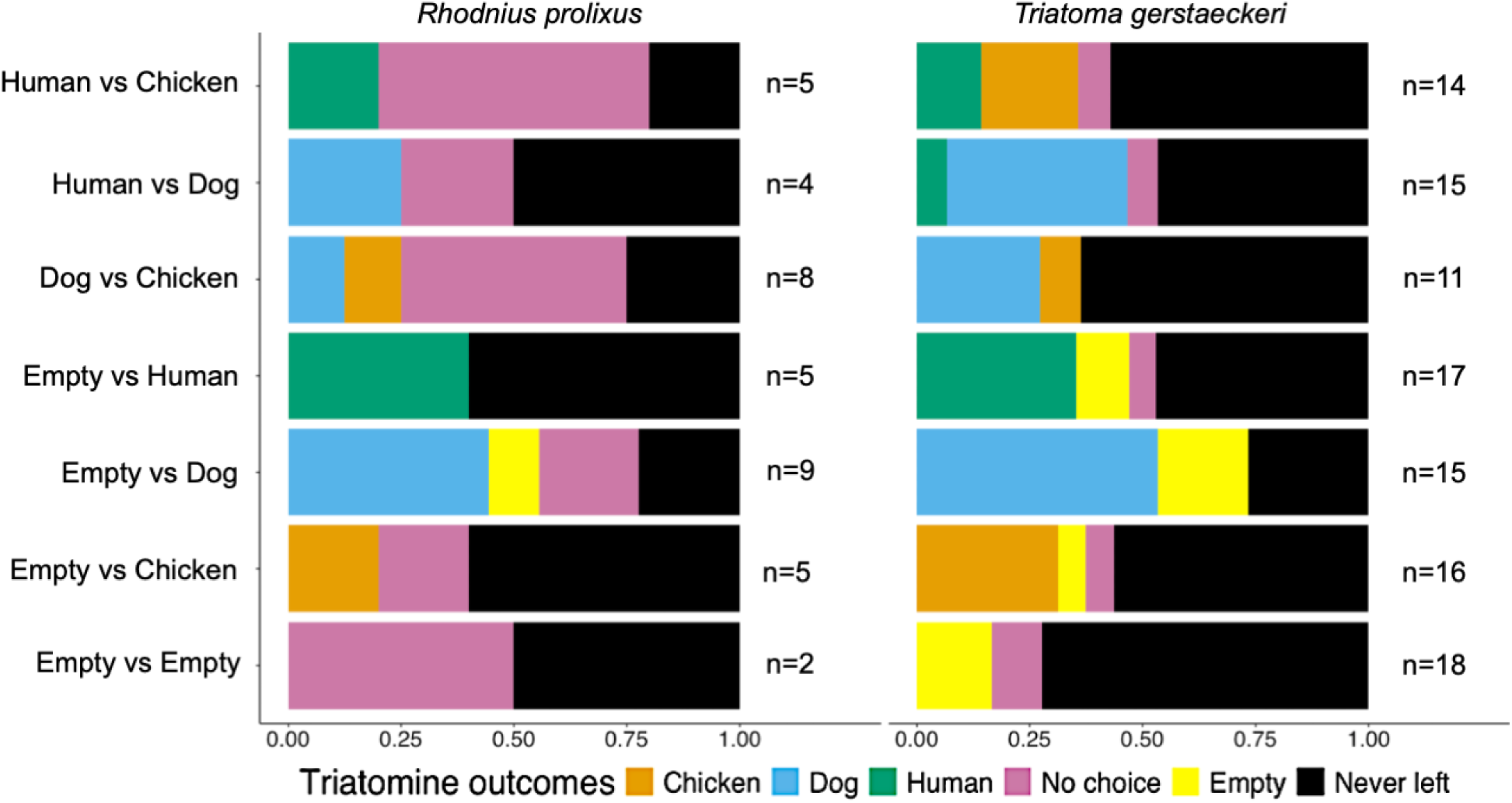
The proportion of trial outcomes among uninfected *T. gerstaeckeri* or *R. prolixus*. Each bar represents the trial combinations, and the colors represent a different outcome. The black numbers to the right of each bar represent the total number of triatomines used for that trial combination.

During the trials, exclusively red lights were used in the room [43] (Figure 1). A YI Lite Action camera (v1. 8.21, YI Technology, Pudong District, Shanghai, China) was placed above the central connecting tubing to record the triatomine activity (Figure 1). A series of abiotic measurements were recorded in three locations: within the room and on both sides of the central connection tube: temperature °C, humidity %, CO_2_ ppm (Amprobe CO_2_-100 handheld CO_2_ meter, Everett, Washington and TEKCOPLUS CO_2_ monitor, Kowloon, Hong Kong, China) and wind speed m/s (Testo handheld anemometer, West Chester, PA, US). Prior to host and triatomine entrance in the olfactometer, baseline measurements were recorded and wind speed on both sides of the chamber were ensured to be equal within 0.3 m/s of each other. Once hosts were enclosed into the chambers, the timer of the trial started. After 5 min, the same abiotic measurements were recorded again and one individual triatomine was introduced into the starting point of the olfactometer (Figure 1). The abiotic measurements were repeated every ten minutes for the full trial. Additionally, temperature and humidity maximum and minimum were recorded for the duration of the trial within the host chambers. Initially, trial duration was at maximum 60 minutes, however this time was reduced to a maximum of 40 min after 58 trials, as we observed most triatomines who moved during the trial initiated movement within the first 10 min and 45 s and making decisions within the first 22 min and 17 s on average. A trial ended when one of the following occured: 1) a choice is made by the triatomine as defined by falling into the terminal pitfall trap (Figure 1); 2) the full trial time has elapsed; or 3) a host displays signs of discomfort that meet the animal use protocol requirements to terminate the trial. This includes any stress signals, such as vocalizing or scratching, displayed by the dog or chicken for more than 15 mins.

In between each trial, the host chambers were cleaned with 10% bleach followed by 70% ethanol, and the central connection tube was replaced with a clean and dried set. We constructed five identical central connecting tube and the starting point (Figure 1) which were washed with 1% Alconox Powdered Precision Cleaner (Alconox, Inc., White Plains, NY, US) solution, then rinsed with 70% ethanol, and allowed to air dry before reuse.

During each trial, the following triatomine behaviors were measured: 1) time for a triatomine to leave the starting point of the olfactometer and begin advancing towards hosts, 2) the side of the olfactometer the triatomine enters (i.e., toward left chamber versus right chamber), 3) the amount of time the triatomine is advancing towards host chambers, and 4) the time at which the triatomine committed to a host choice. Video footage was reviewed to calculate the time (seconds) the triatomines spent in either side of the olfactometer after they had left the starting point. To facilitate this process, a URL software was created to track time spent in an area of a video by overlaying adjustable boxes and then clicking in that box anytime the organism of interest enters that space [44].

Possible outcomes of a triatomine in the olfactometer choice assay included 1) a choice (i.e., dog, chicken, human, empty), 2) no choice, and 3) never left. Triatomines made a choice by crossing the threshold and falling into the pitfall trap at the junction of the clear connecting tube and the host chamber (Figure 1). A “no choice” response means the triatomine left the starting point and advanced towards any side of the olfactometer but made no commitment to a choice by crossing over the threshold to fall into the pitfall trap. The “never left” outcome represented a triatomine that never left the starting point where they were initially placed into the olfactometer. *Statistical analyses*

The statistical analyses can be summarized into four broad research questions: 1) is the dual-choice olfactometer design useful in identifying triatomine foraging behaviors, 2) what influences a triatomine to move towards a host, 3) is *T. cruzi* infection associated with different foraging behaviors, and 4) is triatomine species associated with different foraging behaviors. All statistical analyses were conducted in R Version 1.4.1717 where statistical differences were determined with *P*-values less than 0.05.

To test the dual-choice olfactometer design, we ran separate binomial exact tests (*‘binom.test’* function within the *’stats’* package) [45] for: 1) significant differences between the frequency of choosing the left versus right chamber, 2) if starving triatomines left the starting point when offered a host (the positive controls of any host in a chamber and just one host offered), and 3) if triatomines didn’t leave the starting point during negative controls (no hosts offered in either host chamber). A t-test was used to identify any significant differences between mean seconds spent on either side of the olfactometer design.

To better understand foraging behaviors of starving triatomines in the presence of known associated hosts, we ran generalized linear models with the ‘*lme4*’ and ‘*MASS*’ packages [46,47]. We tested whether a triatomine leaving the starting point (binomial response variable) was associated with predictor variables including *T. cruzi* infection status, number of days the triatomine was starved, triatomine species, the time of day that the trial was conducted, or the humidity and temperature of the experimental room.

With generalized linear models, we tested if *T. cruzi* infection status (binomial response variable) had an association with any of the trial outcomes (i.e., never left, no choice, empty, dog, human, or chicken) among *T. gerstaeckeri*. Furthermore, among the *T. gerstaeckeri* that left the starting point, we tested separately if there was a significant association between *T. cruzi* infection status (binomial response variable) and predictor variables such as i) if the triatomine moved into both sides of the olfactometer, ii) the seconds a triatomine spent near each outcome (i.e., host or empty), and iii) the seconds they took to begin moving towards the host options. Odds ratios were calculated from each generalized linear models identifying associations between *T. cruzi* infection status and the trial outcomes of triatomines that left the starting point.

To test differences among *T. gerstaeckeri* and *R. prolixus*, we used a generalized linear model to identify if either of these species (binomial response variable) had significant associations with the different trial outcomes (i.e., never left, no choice, empty, dog, human, or chicken). Similarly, as previous models, we also tested for species associations and spending time near hosts, the time they took to begin advancing towards hosts, the time they took advancing towards hosts, and the time they spent near each outcome. Odds ratios were calculated from each generalized linear models, identifying associations between triatomine species and the trial outcomes of triatomines that left the starting point.

In attempt to identify which of the abiotic outputs of each host type is associated with host choice, we ran separate generalized linear models for each host type. Predictor variables included abiotic outputs measured in the host chamber (average CO_2_ (ppm), temperature (°C), and relative humidity (%)), host height (cm) and weight (kg) and the outcome was a binomial response variable (whether or not the host was chosen). Odds ratios were calculated from each generalized linear models.

Kruskal-Wallis rank sums tests were used to identify significant differences between mean days starved among all triatomines that made a host choice and between mean abiotic outputs of each host type. Pearson’s Chi-squared tests were used to test for differences in triatomine exploration of both sides of the olfactometer depending on *T. cruzi* infection status or species.

## Results

### Overall triatomine and host descriptions

A total of 238 trials with individual triatomines were conducted between October 2021 and August 2023. We used 225 trials in the analysis and excluded 13 trials due to host discomfort or olfactometer malfunction (Table 1). Starvation of triatomines ranged from 13 to 170 days, with an average of 44 days. There were 187 *T. gerstaeckeri* and 38 *R. prolixus* used in the trials. Among the *T. gerstaeckeri*, one was a 4^th^ instar, 156 were 5^th^ instars, 26 were adults, and four did not have their life stage recorded. Among the *R. prolixus*, 23 were 5^th^ instars, 13 were recently molted adults, and two did not have their life stage recorded. This sample size included 81 *T. cruzi-*infected *T. gerstaeckeri* which all had a qPCR positive fecal spot before their trial.

**Table 1:**
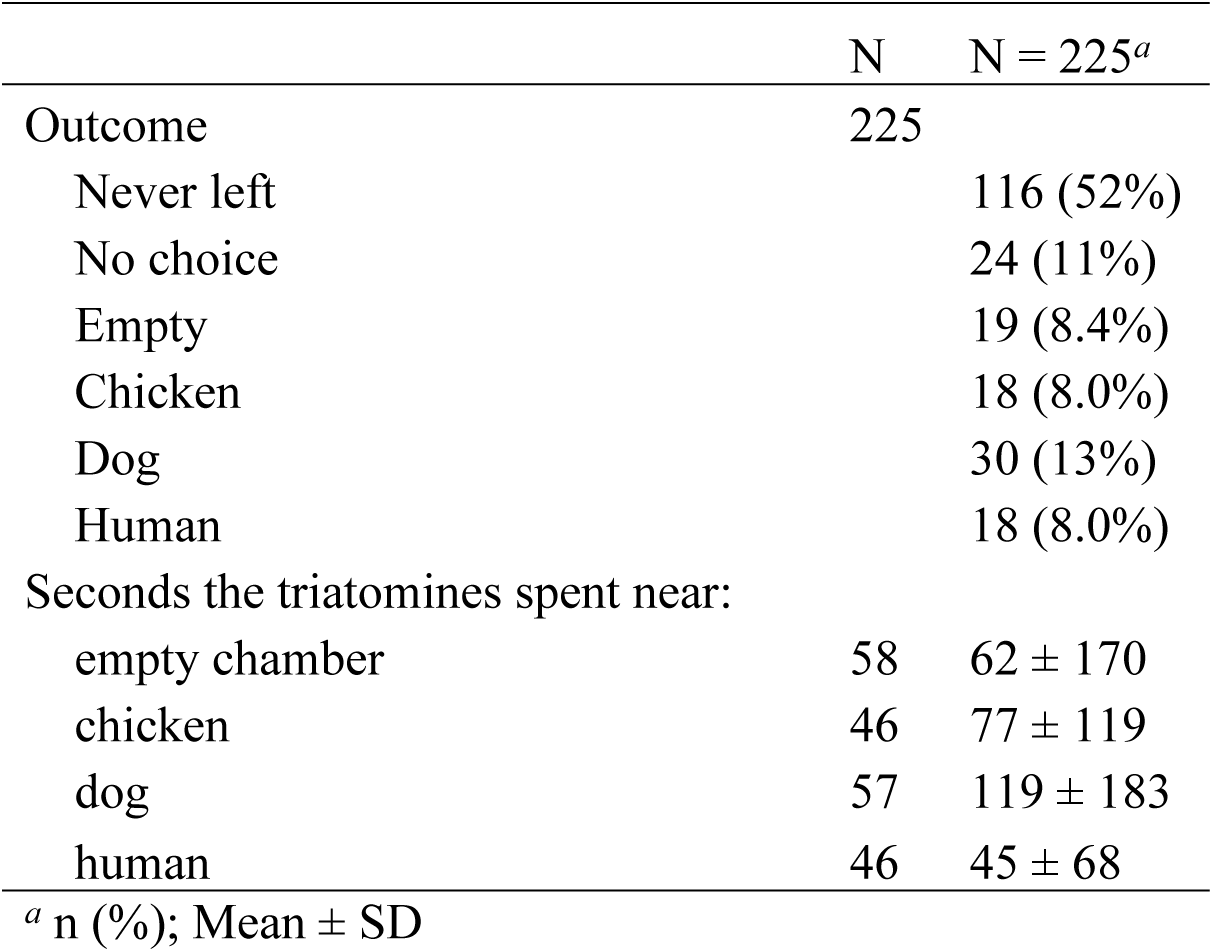
Summary of trial outcomes and the mean seconds spent near a host chamber across trials with *Triatoma gerstaeckeri* and *Rhodnius prolixus*.

Fifteen dogs were evaluated for the study; ten were used in one or more trials each (total of 102 trials with dogs) and five dogs were excluded due to either incompatibility with sitting in a box because of stress, or they had a positive *T. cruzi* screening result (see Supplemental Materials). Among the ten dogs used in trials, the average weight was 117.3 kg (ranging from 24.26 kg to 205.1 kg), the average height 54.4 cm (ranging from 32 cm to 64.14 cm), and the average age was 6.5 years (ranging from 3 to 12 years). The measurements from chambers with dogs included average temperature of 25.05 °C (SE± 0.92), an average humidity of 50.5% (SE± 2.43), and an average of 1,222.13 ppm CO_2_ (SE± 82.87) during trials.

Twenty humans were enrolled and participated in 93 trials. Average heights of humans were 170 cm (ranging from 152.4 cm to 185.42 cm) and average weight was 76.6 kg (ranging from 61.2 kg to 113.4 kg). The measurements from chambers with humans included average temperature of 24.2 °C (SE± 0.29), 51.3% humidity (SE± 2.25), and 1,620.64 ppm CO_2_ (SE± 110.79) during trials.

Twenty-six individual chickens were used in 99 trials. Chickens were approximately 18-24 weeks old with the average weight of 1.34 kg. The measurements from chambers with chickens included average temperature of 26.3 °C (SE ± 82.87), 44.21% (SE ± 2.1) humidity, and 597.4 ppm CO_2_ (SE ± 55.12) during trials.

### Dual-choice olfactometer design and testing the effect of positive and negative control trials

Among all the choices made (i.e. insect dropped into pitfall trap) regardless of the chamber contents (host or empty), 48 triatomines chose the left chamber and 38 chose the right chamber (*P*-value = 0.3, Binomial exact test). Aggregate data from all trials show triatomines spent on average 79 seconds (± 148 seconds) on the left side and 75 seconds (± 175 seconds) on the right side (*P*-value = 0.9, t-test). In all trials where at least one host was offered (n=197), triatomines left the starting point 51.3% (n=101) of the time. Among the 28 ‘negative control’ trials in which no host was offered on either side, we found that 32% (9/28) of the triatomines left the starting point, including 28% (5/18) of the *T. cruzi*-infected *T. gerstaeckeri,* 65% (5/8) of the uninfected *T. gerstaeckeri,* and 50% (1/2) of *R. prolixus*.

### Overall triatomine trial outcomes

Of the 225 total trials, just over half of the triatomines never left the starting point in the olfactometer (52%, n = 116; Table 1). Among those that did leave the starting point (Table 1), they spent the most time near dogs (119 ± 183 s) and chose dog the most out of the hosts chosen (13%, 30/225). Triatomines that left the starting point did so on average 640 s after being placed into the olfactometer, and the triatomines that committed to a choice (i.e., fell into the pitfall trap) took an average of 786 seconds after being placed into the olfactometer.

### Factors associated with triatomines leaving the starting point to advance towards hosts

From a generalized linear model, we found that the abiotic conditions of the experimental room, the time of day the trial was conducted, and *T. cruzi* infection status were not associated with whether a triatomine left the starting point of the olfactometer. We observed a significant positive association between the number of days that the triatomines were starved and if they left the starting point (*P*-value = 0.03, GLM); triatomines that left the starting point were starved on average 49 d, whereas the individuals that did not leave the starting point were starved on average 41 d. For every additional day the triatomines were starved, they were 1% more likely to leave the starting point (OR:1.02, CI: 1.01 to 1.04). Triatomine species was also significantly associated with leaving the starting point in the olfactometer in which *T. gerstaeckeri* was less likely to leave than compared to *R. prolixus* (OR: 0.13, CI 0.03-0.48). Among all 66 triatomines that made a choice (n=66), the mean number of day starved was 45 ± 21 days, with no significant differences among those that chose chicken (n=18; 50 ± 26 days), dog (n=30; 42 ± 14 days) or human (n=18; 45 ± 24 days); p = 0.9.

### Impact of T. cruzi infection status on host preference and foraging behaviors of T. gerstaeckeri

*Trypanosoma cruzi-*infected *T. gerstaeckeri* had a significant negative association with choosing dogs (*P*-value= 0.05, GLM). Among the *T. gerstaeckeri* that left the starting point, the odds of a *T. cruzi*-infected *T. gerstaeckeri* choosing dogs was 59% (OR: 0.41, CI 0.16- 0.95) less than uninfected *T. gerstaeckeri* (Figure 5). Although we found no significance among the other trial outcomes *T. gerstaeckeri* could have made if they left the starting point, the odds of *T. cruzi*-infected *T. gerstaeckeri* choosing chickens was 22% (OR: 0.78, CI 0.28 - 2.09), humans was 33% (OR: 0.67, CI 0.22 - 1.85), and the empty chamber was 44% (OR: 1, CI 0.39 - 2.56) less compared to uninfected *T. gerstaeckeri* (Figure 5). There was no odds difference between *T. cruzi*-infected and uninfected *T. gerstaeckeri* leaving the starting point but not committing to a choice. We found a higher proportion of uninfected *T. gerstaeckeri* chose dog (16%; 17/106) compared to *T. cruzi-*infected *T. gerstaeckeri* (8.6%; 7/81) (Figure 2).

**Figure 5:**
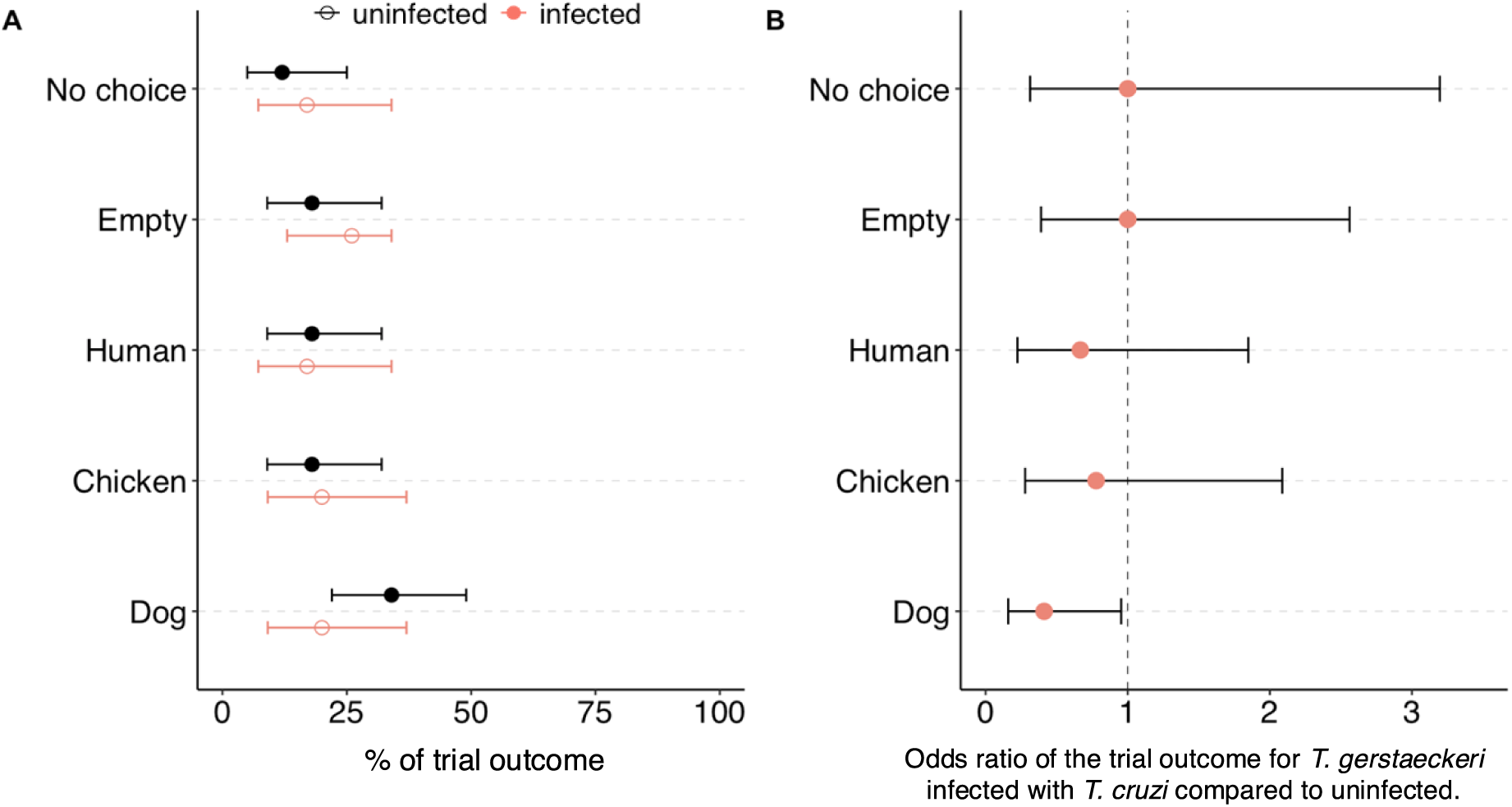
The percentage of trial outcomes of host choice between infected vs. uninfected *T. gerstaeckeri* (A) and the odds ratio of those outcomes (B). The dashed vertical line at 1 indicates the line of no effect.

We found no significant association between *T. cruzi* infection status and if the triatomines spent more time near each of the host chambers (i.e., dog, chicken, human, or empty), if the triatomines explored both sides of the olfactometer, or the time they spent moving throughout the olfactometer. *Trypanosoma cruzi-*infected *T. gerstaeckeri* took less time to begin moving towards host chambers (*P*-value = 0.03, GLM; Figure 3) in which *T. cruzi-*infected *T. gerstaeckeri* took 522 s (± 577 s) on average, whereas uninfected *T. gerstaeckeri* took 801 s (± 704 s) on average. For every additional second that it took for *T. gerstaeckeri* to leave the starting point and advance towards hosts, the odds of the triatomine being *T. cruzi*-infected was 0.06% (OR: 0.9994, CI: 0.9989 - 0.9999) less likely than an uninfected *T. gerstaeckeri*.

### Triatomine species impact on host preference and foraging behaviors

We identified that *T. gerstaeckeri* were significantly associated with choosing dogs (*P*-value = 0.03, GLM), choosing the empty chamber (*P*-value = 0.04, GLM), or never leaving the starting point (*P*-value = < 0.0001, GLM) compared to *R. prolixus* (Figure 4). The odds of *T. gerstaeckeri* choosing a dog was 183% (OR: 2.83, CI: 1.18 - 7.9), choosing the empty chamber was 800% (OR: 9, CI: 1.69 – 165.98), and never leaving the starting point was 300% (OR: 4, CI: 2.29 - 7.47) greater than *R. prolixus* (Figure 6).

**Figure 6:**
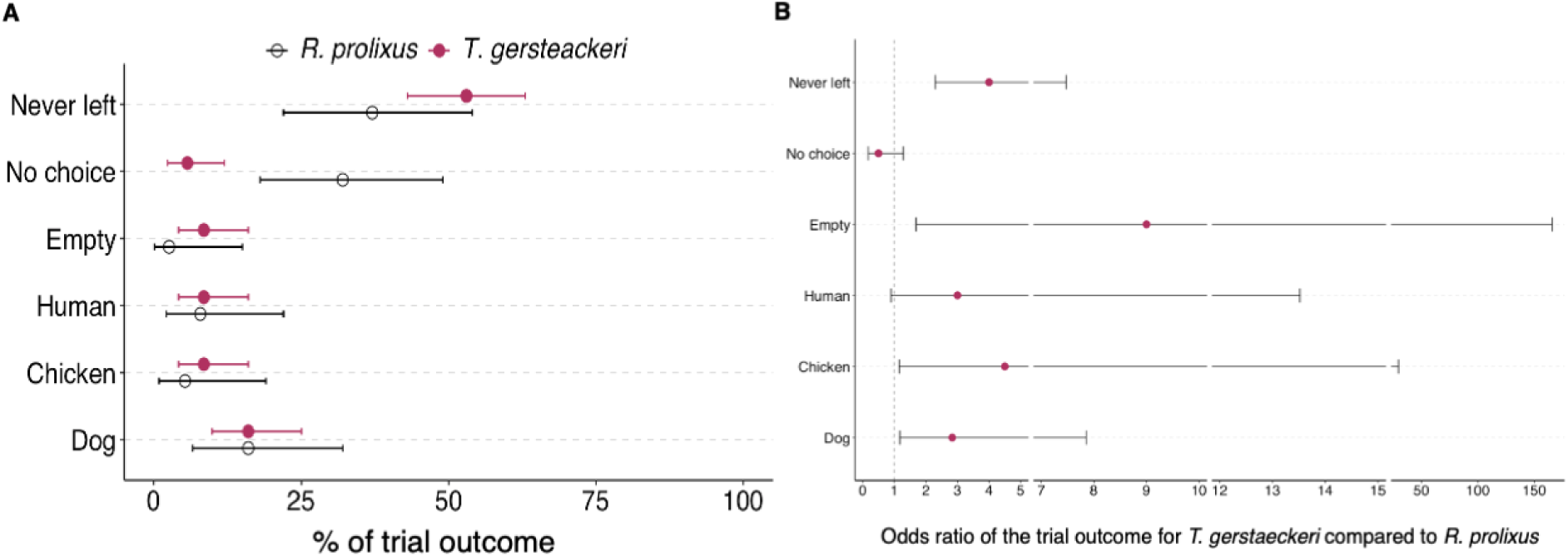
The percentage of trial outcomes of host choice between uninfected *T. gerstaeckeri* vs. *R. prolixus* (A) and the odds ratio of those outcomes (B). There are 3 breaks in the x-axis of B. The dashed vertical line at 1 indicates the line of no effect.

We found *T. gerstaeckeri* were significantly associated with taking longer to begin advancing towards hosts compared to *R. prolixus* (*P*-value = 0.003, GLM), in which *R. prolixus* began advancing towards hosts at 491 s (± 477 s) on average, while *T. gerstaeckeri* took double that time to begin advancing towards hosts (801 ± 704 s) on average. For every additional second that it took for triatomines to leave the starting point and advance towards hosts, the odds of the triatomine being *T. gerstaeckeri* was 0.1% (OR: 1.00102, CI: 1.000413 - 1.001771) more likely than a *R. prolixus* (Figure 6). There was no significant difference between species and the time individuals spent advancing towards hosts or time spent near the host options. *Rhodnius prolixus* explored both sides of the olfactometer (46%, n=11/24) more often compared to *T. gerstaeckeri* (14%, n= 7/50) (*P*-value = 0.003, Pearson’s Chi-squared test).

### Impact of host abiotic cues on triatomines choice

Average temperature and humidity emitted from each host type were not significantly different, while CO_2_ was significantly different (Table 2). Host size and average abiotic host cues (CO_2_, humidity, or temperature) were not significantly associated with triatomines choosing chickens or humans. However, increasing humidity was significantly associated with triatomines choosing dogs (*P*-value = 0.03, GLM), where every percent increase in the humidity level when dogs were a the host chamber, triatomines were 3.81 times more likely to choose the dog (OR: 1.04, CI 1.01 -1.08).

**Table 2:** Summary of the mean abiotic cues emitted from each host type across all trials according to host choice.

| | Number of trials | Chicken<br>N = 42<sup>a</sup> | Dog<br>N = 84<sup>a</sup> | Human<br>N = 54<sup>a</sup> | $P$-value<sup>b</sup> |
| --- | --- | --- | --- | --- | --- |
| Mean Temperature °C | 93 | 26.58 ± 9.33 | 23.68 ± 2.62 | 23.43 ± 3.22 | >0.9 |
| Mean Humidity % | 92 | 51 ± 15 | 55 ± 17 | 51 ± 15 | 0.7 |
| Mean CO<sub>2</sub> ppm | 93 | 756 ± 615 | 1,202 ± 620 | 1,559 ± 893 | **0.002** |

<sup>a</sup> Mean ± SD
<sup>b</sup> Kruskal-Wallis rank sum test

## Discussion

Our data contribute to the growing evidence that *T. cruzi* influences triatomine foraging behaviors which could regulate transmission dynamics [4,23,24,48–52]. Contrary to our hypothesis, *T. cruzi-*infected triatomines did not display a preference for mammalian reservoir hosts over *T. cruzi* incompetent chickens. However, our finding that the *T. cruzi-*infected triatomines showed no distinct host preference and initiated movement towards hosts faster indicates they may be more opportunistic [23,24]. As predation of *T. cruzi*-infected triatomines is a major route of transmission among dogs and wildlife, these changes in foraging behaviors that we observed- namely increased initiation of movement and lack of a host preference- could lead to an increase in vector-host encounters and thus increased opportunities for transmission [3,4]. As previously shown for *Triatoma infestans* [53], we found that uninfected *T. gerstaeckeri* have a preference towards dogs (Table 1, Figure 2 & 5). Given the repeated observations that multiple triatomine species have a preference for dogs; triatomine feeding patterns as measured through bloodmeal analysis indicate high triatomine encounter rates with dogs [18]; and dogs are commonly infected with *T. cruzi* [12], dogs should be a high priority candidate for host-targeted interventions to reduce the triatomines population [32–34].

As expected, triatomines that were starved longer were more likely to advance towards hosts; however, the length of starvation did not impact the triatomines host choice. Variation in the days of starvation was a logistical artifact of our study design, yet allowed us to quantify its impact on foraging behaviors. Although other studies have a drastic variation in days starved before trials [22,24,53], future studies should further investigate this dynamic by controlling the number of days starved and previous bloodmeal host type to investigate the intersection of nutrition, *T. cruzi* infection status, and triatomine foraging behaviors.

Our study illustrates behavioral differences between two different species of triatomines in the presence of known bloodmeals, as we found *R. prolixus* more quickly advanced toward hosts and walked around more within the olfactometer as compared to *T. gerstaeckeri.* Such behaviors may contribute to the parameters of the equation of vectorial capacities for *T. cruzi* [54], potentially increasing vector-host contact.

Importantly, we show for the first time with *T. gerstaeckeri* behavioral differences between *T. cruzi*-infected vs. uninfected individuals in the presence of hosts cues, as has been shown with other triatomine species [23,24,49,51,52,55]. In nature, *T. cruzi-*infected triatomines have different host feeding patterns than uninfected triatomines [18,56]. For instance, a study found that *T. cruzi*-negative triatomines had a higher diversity of bloodmeals compared to the *T. cruzi*-positive triatomines [57]. Differences in host cue olfactory organs and the locomotion gene expression is different between *Trypanosoma-*infected and uninfected triatomines [22,23], potentially driving such observations. Furthermore, it has been documented that *T. cruzi* manipulates triatomine behaviors by increasing their risk of predation by a *T. cruzi* potential host [3,4,21]. Considering these studies in concert with our new results that *T. cruzi*-infected triatomines were faster to respond or advance to hosts and had no particular host preference (Figure 2), *T. cruzi* appears to be associated with vector behaviors that enhance either the insects ability to encounter a host or a hosts ability to detect the insect.

In addition to behavioral differences among wild triatomine species, the time since populations have been colonized compared to wild triatomines can potentially impact foraging behaviors. However, studies show phenotypic plasticity among triatomine mouth parts morphology when fed on different bloodmeal hosts [58,59]; therefore, their behaviors towards hosts may also be adaptable. We included trials using *R. prolixus* from a colony-reared population that started in the 1980’s [60]. We found significant differences in foraging activity between *R. prolixus*- a long-maintained species in a colony and also known as a more anthropogenic species of triatomine [61]- and *T. gerstaeckeri-* a more recently colonized and relatively less anthropogenic triatomine species [62]. We found *R. prolixus* were active more often on both sides of the olfactometer, were faster at advancing towards hosts, and had different host preferences (Figure 4) compared to *T. gerstaeckeri*. The degree to which the observed behaviors reflect generations in colony vs. intrinsic species differences [63], and how the behaviors may scale up to explain differences in the epidemiology of Chagas disease in the regions where each insect species is endemic remains unknown, yet foraging behavior is a critical component in the reproductive rate of *T. cruzi* (*R_0_*) [64].

The ‘experimental cafeteria’ framework allows testing of parasite-vector-host interactions and the manipulation of scenarios to test hypotheses surrounding vector foraging behaviors [53,65], across multiple vector taxa. For instance, the brown dog tick, *Rhipicephalus sanguineus*, preferred humans over dogs at increased temperatures [66], an important finding given changing environmental temperatures. In contrast, we did not find the host size, temperature, or CO_2_ were associated with triatomine choice, but increased humidity was significantly associated with triatomines choosing dogs, suggesting that abiotic factors can influence host choice. In a different olfactometer study. The anthropogenic mosquito *Aedes albopictus* (Diptera: Culicidae) did not prefer human odors in an olfactometer design [68], suggesting that observed human-mosquito feeding patterns are likely due to host availability and not preference. Within the Chagas disease system, a choice study identified that *Triatoma infestans* has a preference for dogs [53], supporting observational bloodmeal analysis studies that commonly reveal dogs are highly utilized host species. In Texas, US, *T. gerstaeckeri* also commonly feed on dogs, but without dog abundance estimates, it is unknown if this pattern reflects a higher availability of dogs or because triatomines preferred to feed on them [18,69].

Limitations of our study include that we reused some host individuals across multiple trials. Our initial study was not designed to uncover within-host factors that contribute to host choice, yet such factors must be considered in larger experimental trials. Lastly, we detected differences in behaviors between *T. cruzi*-infected and uninfected triatomines; however, we only infected triatomines with *T. cruzi* DTU TcI. Considering there are seven different DTUs of *T. cruzi* and they all have different epidemiological affiliations, it is likely that they interact with the triatomine immune system differently and therefore cause different behavioral outcomes [70,71]. Further investigations should test how different *T. cruzi* DTU’s may manipulate foraging behaviors of triatomines differently or how the hosts *T. cruzi* infection status may alter the foraging responses of triatomines [5].

Data from previous studies indicate *T. cruzi* infections impact triatomine behavior [4] and fitness [70] in ways that can influence transmission dynamics. For instance, *T. cruzi-*infected triatomines fly further, have a higher probing or biting frequency, can orientate to hosts faster, defecate faster, and are more likely to approach humans [49,51,52,55,72]. With this experimental design, we were able to detect that *T. cruzi* is also associated with altered host preference. As vector-borne diseases continue to be a global issue, an understanding of the evolutionary and ecological interactions between parasites, vectors, and host can provide foundational information to develop disease control. Potential intervention strategies that can be extrapolated from these results include building triatomine traps via synthetic pheromones of preferred hosts [67,73] and targeting key preferred host species with systemic insecticides [30,33]. Further research should build on this experimental design to test other species of triatomines, other host species, and manipulate the environment to test interactions between *T. cruzi*, triatomines, and hosts.

## Acknowledgments

A special thank you to the dog volunteers (named with permission): AJ, Cinder, Dakota, Doris, Frank, Freya, Gonzo, Madigan, Millie, Onyx, Pana, Sadie, Tally, Tofu, and Yuki. We would like to acknowledge all the human volunteers for sitting in the olfactometer boxes and Koyle Knape for maintaining the chickens. We thank Cassandra Durden, Lisa Auckland, Haydee Montemayor, Lauren Cole, Rachel Busselman, Ed Davila, and Juan Pablo Fimbres-Macias for assistance in data collection and permitting.

